# Early Prenatal Cell-Free DNA Methylation Landscape Informs Epigenetic Purification of the Circulating Fetal Genome

**DOI:** 10.64898/2026.10.20.760549

**Authors:** Tiepeng Liao, Chinmay Rao, Siya Iyer, Spencer C. Ding, Jingru Yu, Zackary Lui, Cindy Yang, Wei Gu

## Abstract

Circulating fetal cfDNA is the analyte of a screening test used by millions of pregnant women annually. However, low fetal fraction limits current testing. Fetal cfDNA can be enriched via hypomethylation, but the informative sites require a first-trimester fetal methylome reference. Here we sequenced fetal cfDNA (median purity 94.7%) derived from discarded media from cultured chorionic villus biopsies (n=30) and assembled a first-trimester methylome across 26M CpG methylation sites at a median depth of 207x. We then built an analytical enrichment model that identified millions of CpG markers and assigned predicted enrichment scores. Guided by this model, we adapted epiSIFT as a genome-wide assay selective for unmethylated DNA molecules. In 28 prenatal plasma cases, epiSIFT raised the median fetal fraction from 5.1% to 28.4% (5.6-fold gain) using the top eighth of markers. These findings establish a first-trimester fetal cfDNA methylome reference and demonstrate epigenetic enrichment of the fetal genome.

## Introduction

Each year, millions of pregnant patients are screened for the common fetal aneuploidies (trisomies 21, 18, and 13) by non-invasive prenatal testing (NIPT), which analyzes fetal-derived placental cell-free DNA (cfDNA) in the maternal circulation. NIPT is highly accurate [1,2], and professional societies like the American College of Obstetricians and Gynecologists (ACOG) now recommend offering it to all pregnant women as the most sensitive and specific screen for these aneuploidies.

Low fetal fraction in the first trimester constrains both NIPT and emerging non-invasive prenatal diagnosis of monogenic disease. First, ~3-5% of NIPT results for aneuploidy are non-reportable, most often because the fetal fraction falls below the ~4% assay threshold [3]. Second, low fetal fraction blocks low-cost platforms that can broaden access. NIPT remains confined to centralized laboratories that run large, capital-intensive sequencers and reliable transportation infrastructure, which is lacking in many regions of the world. Cheaper, rapid alternatives such as limited sequencing panels and digital PCR are being explored [4], but work reliably only at higher fetal fractions, as in second-trimester amniotic fluid [5]. Third, early prenatal detection of monogenic diseases is increasingly important given emerging gene-editing therapies, and non-invasive detection improves dramatically with fetal fraction [6–8]. Together, these observations suggest that amplifying the fetal fraction (i.e. to >30% in the first trimester) could extend the reach of non-invasive screening. To date, no practical method enriches first-trimester fetal cfDNA to purities >30%.

We hypothesized that fetal cfDNA could be enriched, or background maternal cfDNA depleted, by exploiting methylation differences between fetal placental trophoblast-derived cfDNA and background cfDNA, namely the global hypomethylation of trophoblasts [9,10]. Covering this genome-wide requires two comprehensive methylome maps: one of normal plasma, as recently reported by Loyfer et al. [11], and one of the target fetal cfDNA. To our knowledge, the latter does not exist for early pregnancy (9–13 weeks’ gestation), the window most relevant to NIPT. Such a reference would need breadth across the ~28M CpG sites in the human genome and the depth required for precise, nucleotide-resolution methylation measurements.

We therefore generated a fetal reference set from cfDNA in the culture media of chorionic villus (placental) biopsies and confirmed its fetal origin and purity. Combining this with existing normal plasma references, we modeled and mapped the genome-wide potential for fetal enrichment based on methylation differences. We then used a genome-wide epigenetic purification assay [12], epiSIFT (Epigenetic Signal Increase by Filtering for Targets), to enrich fetal cfDNA, demonstrating its efficacy in titrations and in a cohort of 28 NIPT cases.

## Results

### Establishing a First-Trimester Fetal cfDNA Methylome Atlas

To establish a comprehensive, first-trimester fetal cfDNA methylome atlas at base resolution, we used residual media from chorionic villus sampling (CVS), which is a placental biopsy in the first trimester (Fig. 1a). These media are routinely discarded during clinical use and contain high-purity fetal cfDNA from the removed placental syncytiotrophoblast cells.

**Figure 1:**
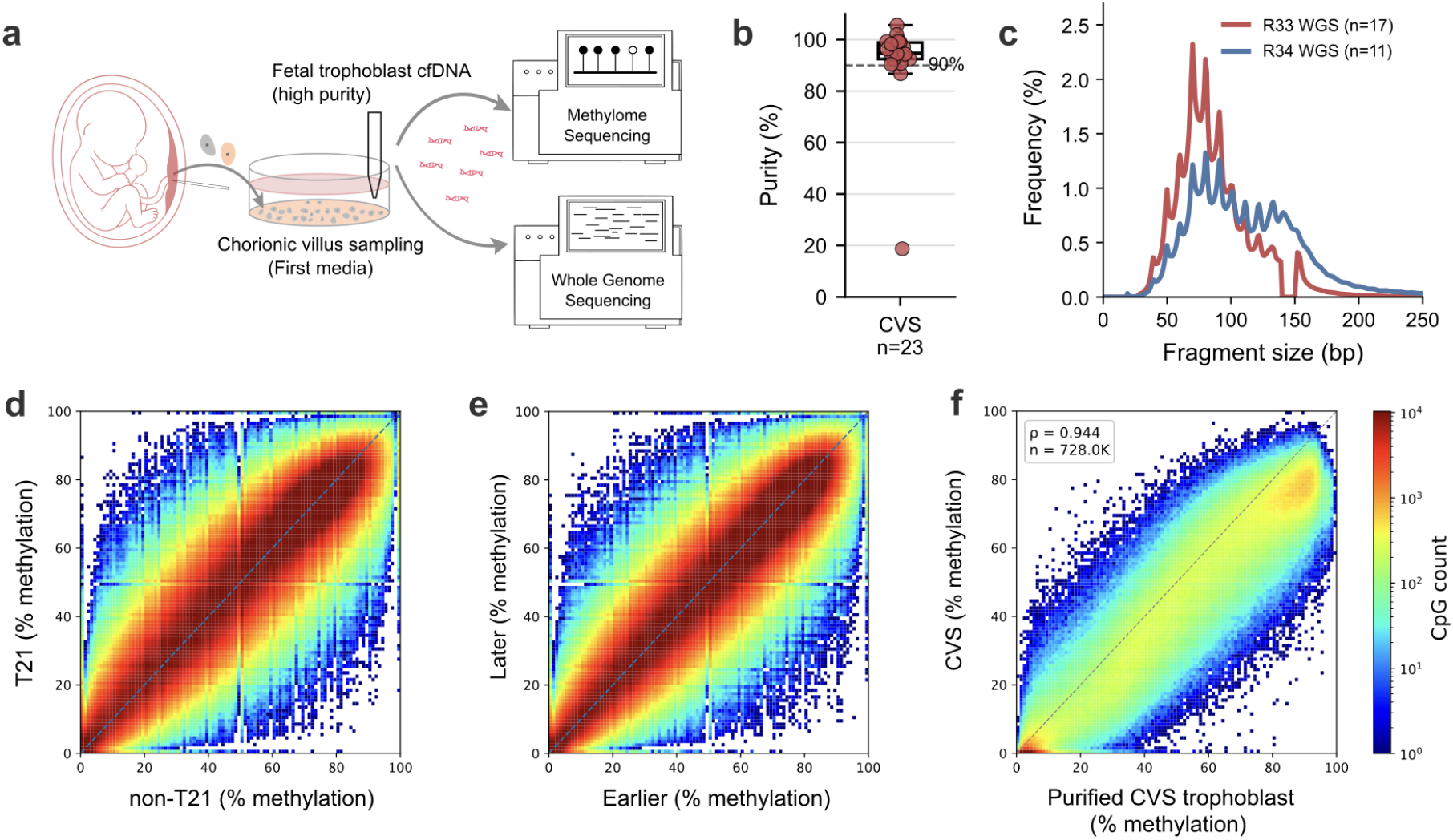
Establishing a First-Trimester Fetal cfDNA Methylome Reference. **a**, Schematic overview of fetal cfDNA recovery from discarded first-trimester CVS culture media, followed by EM-seq for methylome reference construction and whole genome sequencing for purity and fragment-length assessment. **b**, Estimated fetal purity of each CVS sample where purity could be estimated. The dashed line marks 90% purity. We excluded the outlier with lower purity from the final integrated reference. **c**, WGS-derived fragment-length distributions of CVS culture-media cfDNA, shown separately before (n=17, run R33) and after (n=11, run R34) physical size selection. Curves represent averages of normalized frequency distributions across each sample. **d**, CpG methylation concordance between trisomy 21 and normal karyotype CVS samples. Each CpG had a minimum aggregate depth of 10 in both groups. **e**, CpG methylation concordance between CVS samples split into earlier and later gestation age groups, using the minimum depth of 10. **f**, Comparison of the integrated CVS methylome reference with purified placental trophoblast methylation microarray profiles from Yuan et al. For density heatmaps, color indicates the number of CpGs within each bin; CVS, chorionic villus sampling; cfDNA, cell-free DNA; CpG, cytosine-phosphate-guanine; T21, trisomy 21.

We saved CVS culture media from 30 patients, and ultimately included 28 samples in the methylome reference. We excluded one as a holdout sample (see *Enrichment of Trisomy 21 in cfDNA Titrations* section), and another as a potential outlier in purity (see below). These 28 cases had a median gestational age of 12w6d (IQR 12w2d-13w0d) and a median maternal age of 36.5 (IQR 33.75–38.25) years (Supplementary Table 1). Using the extracted cfDNA from these media, we prepared methylome libraries using EM-seq [13] and sequenced to a median depth of 7x (IQR 7–9.75x) across each case, and an integrated reference depth of 207x (IQR 130-306x) across 26,265,480 of approximately 28M CpGs in the human genome.

To verify fetal cfDNA purity, we also created copy number plots from separate shallow whole-genome sequencing (WGS) libraries. We estimated fetal purity in 23 cases using informative trisomy or sex chromosome copy number differences, or the average of the two when available (Fig. 1b) [14]. The median purity was 94.7% (IQR 92.5–98.9%). The one outlier, at 18.7%, may reflect placental mosaicism for monosomy X, but we excluded it out of caution, yielding the 28 cases used in the reference.

We next characterized the length of the fetal cfDNA from culture media with WGS (Fig. 1c, Supplementary Fig. 1). The mean fragment length was 97.6 bp averaged across samples, shorter than previously reported fetal lengths of 143 bp [15]. This may be due to the degradation of fetal cfDNA during incubation at 37 °C and the lack of removal of degraded cfDNA, as would occur in the body. In the latter 11 samples (run R34), we performed physical size selection, yielding longer fragments (Fig. 1c).

Many of the cases involved fetuses with aneuploid genetics due to using CVS as a source. However, all cases were integrated, as we observed a high correlation in the methylation values between Trisomy 21 cases (n=13) and 11 normal karyotype cases (Spearman’s ρ 0.930, Fig. 1d). Similarly, the correlations between the youngest half of all gestational ages and the older half of all gestational ages, as well as for fetal sex, were high at Spearman’s ρ 0.941 and 0.936, respectively (Fig. 1e, Supplementary Fig. 2a).

**Figure 2:**
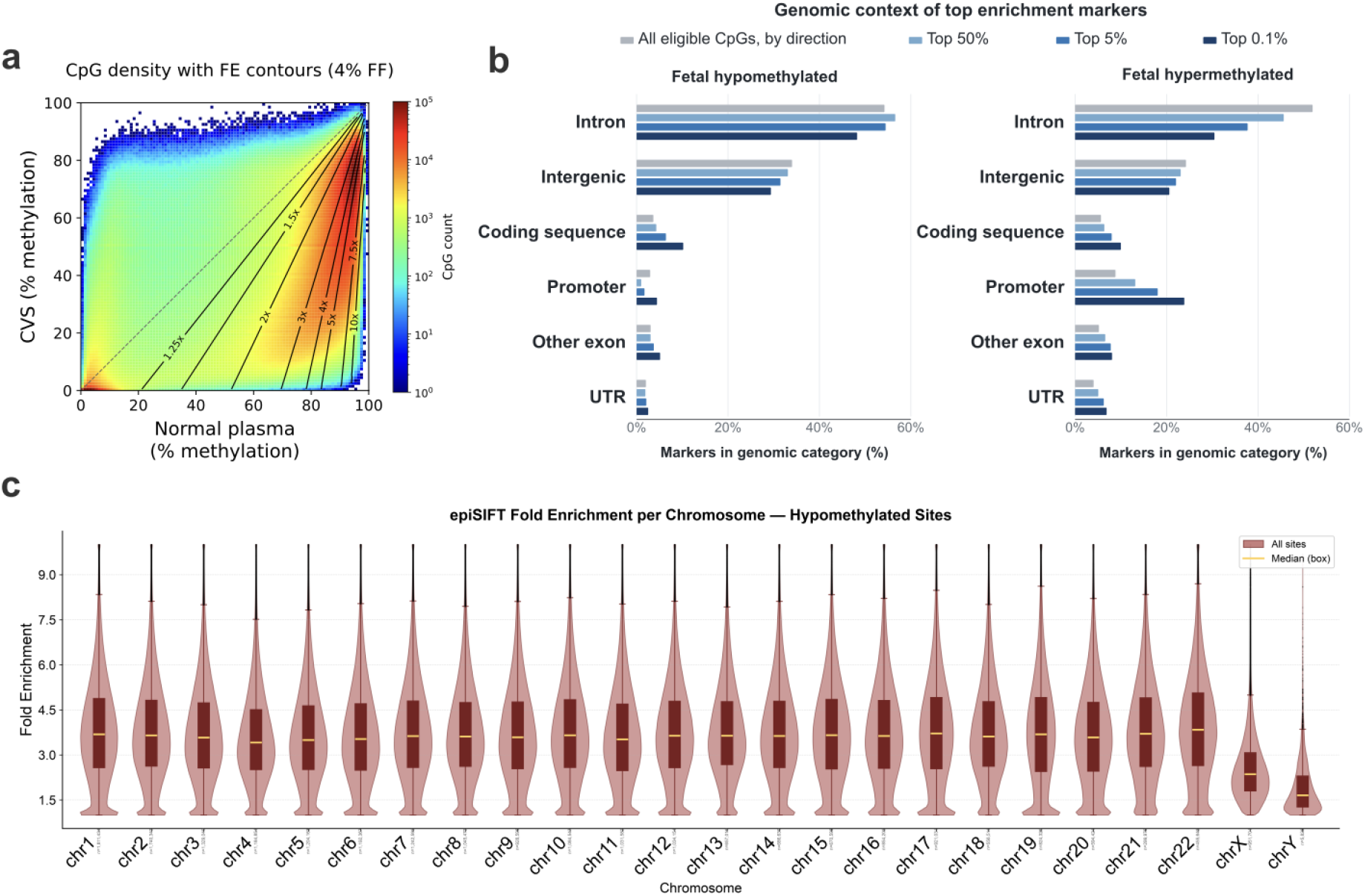
Enrichment Model. **a**, Density heatmap of CpG methylation in the integrated CVS fetal cfDNA and Loyfer et al. normal plasma references, overlaid with theoretical fold-enrichment contours assuming a 4% starting fetal fraction. Minimum depths were 50 for CVS and 450 for normal plasma. Heatmap color indicates the number of CpGs per bin. **b**, Mutually exclusive genomic regions across CpGs and the top 50%, 5%, and 0.1% model enrichment markers, shown separately for hypomethylated and hypermethylated CpG markers. **c**, Violin and boxplots of the chromosomal distribution of theoretical fold enrichment for hypomethylated CpG markers. CVS, chorionic villus sampling; CpG, cytosine-phosphate-guanine; FE, fold enrichment; FF, fetal fraction; UTR, untranslated regions (of exons).

To verify the accuracy of this integrated fetal methylome, we compared it against microarray data (728k CpGs) from Yuan et al., who physically purified trophoblast cells from 9 CVS samples (data from 5 of 9, GEO GSE159526) [9]. A correlation heatmap showed a strong correlation (Spearman’s ρ 0.944) (Fig. 1f). Compared with the purified CVS cells, our reference skews slightly less methylated (hypomethylated), which may reflect greater representation of late-stage syncytiotrophoblastic cells that directly shed fetal cfDNA into maternal circulation. This difference is notable because it reflects the epigenetic phenomenon that underpins our proposed enrichment.

Finally, we assessed whether any individual CVS sample disproportionately influenced the integrated reference by comparing each sample with a reference constructed from the remaining CVS samples. Across the 28 samples included in the reference, all samples remained concordant in a leave-one-out analysis, with the lowest Spearman’s ρ of 0.753 and median Spearman’s ρ of 0.818. This supports the consistency of the integrated fetal methylome and argues against dominance by a single sample (Supplementary Fig. 2b).

### Epigenomic Mapping and Epigenetic Purification of Fetal cfDNA

With the establishment of a high-depth fetal cfDNA methylome atlas, we sought to map the maximal theoretical fetal enrichment across the genome and define the most potent enrichment markers. To do this, we combined the current fetal atlas (the target signal) in Figure 1 with high-depth plasma data from healthy volunteers as background cfDNA [11] to build a unified enrichment model. Because pregnant women are younger than the range of healthy volunteers in the plasma cohort, we correlated methylomes between younger and older volunteers. We found they were highly similar (Spearman’s ρ = 0.965; Supplementary Fig. 3), and therefore integrated data from the entire healthy plasma cohort regardless of age.

**Figure 3:**
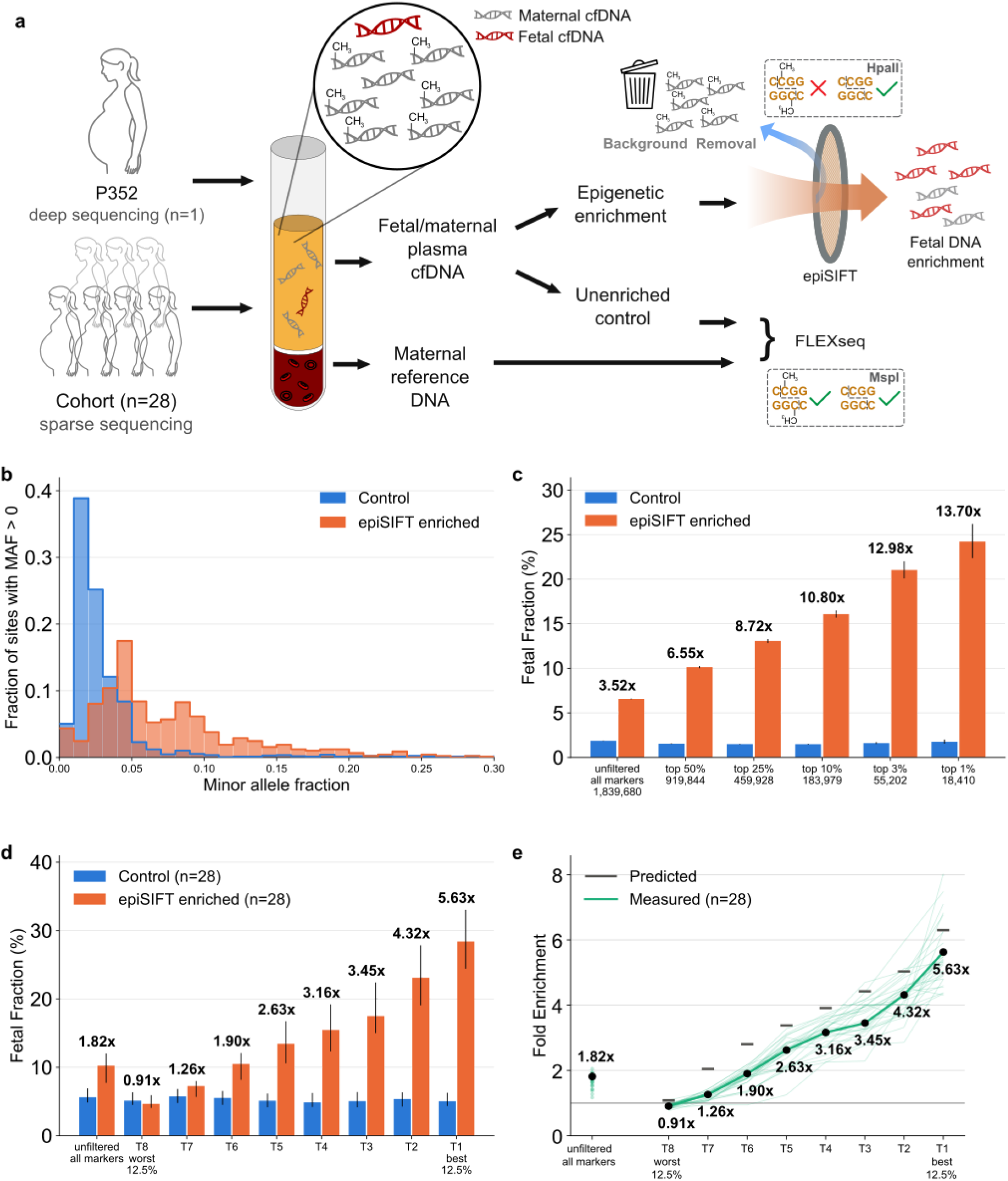
Enrichment for fetal cfDNA in maternal blood plasma. **a**, Schematic illustrating circulating fetal cfDNA in the maternal blood and experimental designs on an index case with deep sequencing (P352) and a cohort of 28 NIPT plasma cases. **b**, Histograms of minor allele fraction (MAF) at maternal SNPs that are homozygous in P352 buffy coat, control versus epiSIFT-enriched, top 10% markers. MAF is the non-maternal allele count divided by the summed depth of the two panel alleles. **c**, Fetal fraction and fold enrichment based on top markers up to 1%, yielding up to 13.7-fold enrichment. **d**, Fetal fraction and fold enrichment in the cohort of 28 cases across 8 tiers (T1-T8) of increasingly potent markers, yielding up to 5.6-fold enrichment. **e**, Predicted and measured fold enrichment across 8 tiers.

We first mapped methylation levels (betas as % methylated at one site) in the target and background references across the human genome (Fig. 2a). Of the approximately 28M CpGs in the human genome, 23,546,534 CpGs had a minimum depth of 50 in the integrated fetal cfDNA reference and 450 in the normal plasma reference. Of note, most CpGs were highly methylated in the plasma, whereas variably methylated in CVS.

We then developed a theoretical model in which methylated molecules were in the background, and unmethylated molecules were the fetal target. We modeled the sample as a mix of fetal target fraction p and background fraction 1-p, where β_*T*_ and β_*B*_ are the methylation fractions at a CpG in the target and background references, respectively. The model can be reduced to Equation 1 to calculate the fold-enrichment (FE) for hypomethylated markers (β_*T*_ < β_*B*_

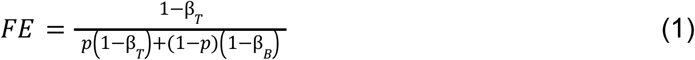

and Equation 2 for hypermethylated target biomarkers (β_*T*_ < β_*B*_):

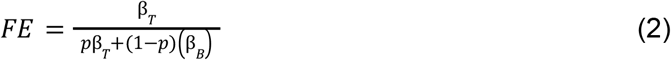

We overlaid our model on top of the target and background CpG density map (Fig. 2a). High-density CpGs are hypermethylated in plasma and hypomethylated in CVS, and 4,623,057 CpGs could theoretically reach >5-fold enrichment from a 4% starting fetal fraction. Most markers fall in intronic and intergenic regions regardless of enrichment potential (Fig. 2b), though the few hypermethylated markers with high enrichment potential skewed toward promoter regions, as did the weakest hypomethylated markers (Supplementary Fig. 4). Since markers were overwhelmingly hypomethylated rather than hypermethylated, we focused exclusively on the hypomethylated marker set.

**Fig. 4:**
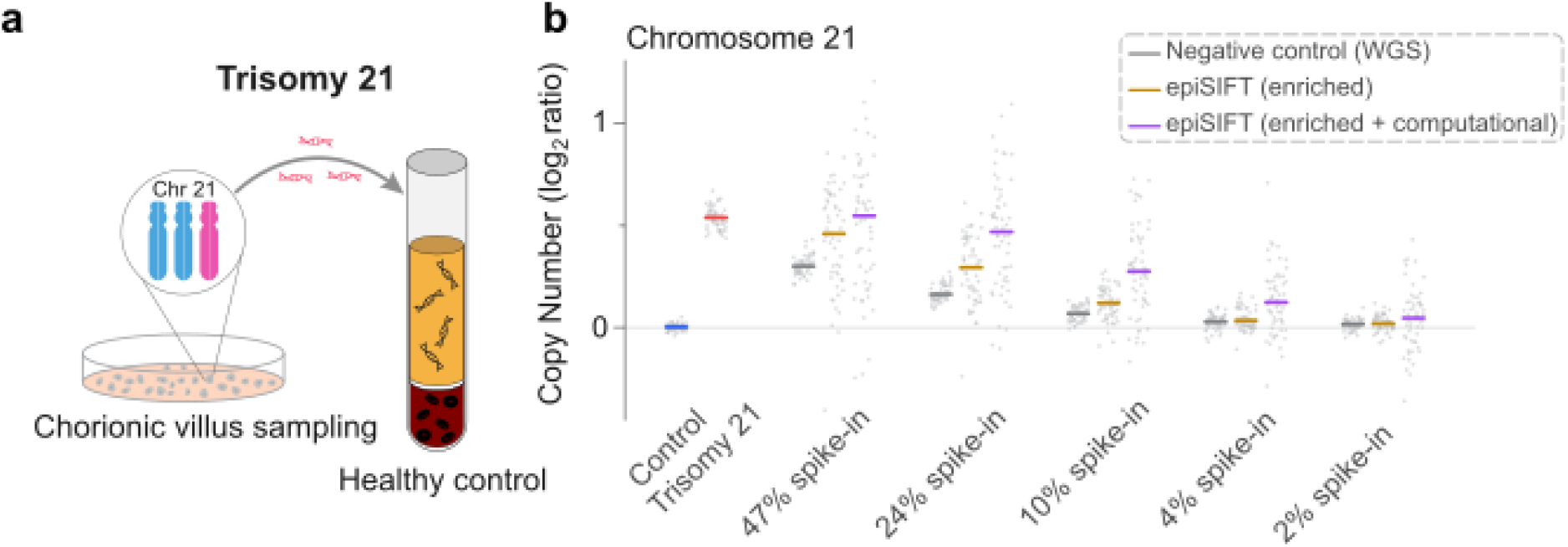
Enrichment for fetal cfDNA with Trisomy 21. **a**, Schematic of positive fetal cfDNA (trisomy 21), derived from culture media of a placental CVS biopsy (case BF4232), spiked into a control plasma cfDNA. **b**, Copy number plots of chromosome 21 across titrations measured at 47, 24, 10, 4 and 2% by control WGS (planned 30, 10, 3, 1 and 0.3%), and the epiSIFT assay. Control (unspiked plasma cfDNA) and Trisomy 21 (undiluted fetal cfDNA) serve as negative and positive controls.

To benchmark our reference against published fetal differentially methylated regions (DMRs), we evaluated 31 CpGs within 11 chromosome 21 DMRs. Two CpGs had absolute methylation differences below 0.05 and were considered neutral. Our fetal and plasma references reproduced the reported fetal hypermethylation at 28 of the remaining 29 informative CpGs [16]. Predicted enrichment ranged widely from 1.00-to 7.06-fold, but only 11 of 31 exceeded 5-fold. Deep sequencing at base resolution revealed differences in predicted enrichment among CpGs within published DMRs, supporting CpG-level marker selection.

More broadly, autosomes had similar enrichment performance, whereas the sex chromosomes performed worse (Fig. 2c). Assuming a 4% starting fetal fraction, the autosomal median fold enrichment among hypomethylated CpGs was 3.61-fold, compared with 2.36-fold for chromosome X.

### EpiSIFT Enrichment Assay and FLEXseq Control Assay

Our fetal methylome mapping and enrichment model provided theoretical evidence that enrichment by methylation can be applied to fetal cfDNA. To achieve this, we hypothesized that we could adapt our previously reported assay, which depletes methylated human background to enrich microbial cfDNA [12]. We term the assay used here epiSIFT (Epigenetic Signal Increase by Filtering for Targets) (Fig. 3a). The assay uses a methylation-sensitive restriction enzyme (HpaII) in combination with adaptors previously reported in our FLEXseq technique [17] to sequence fragments cut only once by the enzyme while excluding uncut fragments, which comprise methylated fragments and fragments not containing CpGs of interest. FLEXseq, which uses the MspI enzyme, can be used as an unenriched control, covering overlapping regions of the genome while equally representing methylated and unmethylated DNA.

### Fetal Enrichment Demonstrated in Maternal Blood Plasma

To first test fetal cfDNA enrichment from the blood plasma of a pregnant woman, we measured fetal fractions with common single-nucleotide polymorphisms (SNPs) enriched by the epiSIFT assay, whereas control libraries used MspI rather than HpaII. We isolated plasma cfDNA and leukocyte DNA for germline analysis (Fig. 3a).

EpiSIFT produces a notable visual shift in the minor allele fraction of maternal homozygous and fetal heterozygous positions as an indicator of increasing fetal fraction (Fig. 3b). To estimate the exact fetal fraction, we used a previously validated binomial-mixture maximum likelihood estimator (MLE) that incorporates the maternal-homozygous (based on maternal cells) SNP allelic counts adjacent to CpG selector markers, the population frequency of those SNPs, and an estimated global error rate using alternative bases at those SNP positions [18,19]. We performed deep sequencing on one index case (P352), yielding 115M unique control (MspI) reads and 63M unique epiSIFT (HpaII) reads. Using all 1.8M predicted markers that overlap with the assay, we found a 3.52-fold enrichment in the index case, whereas the top 1% (18k) of markers yielded a 13.7-fold enrichment (Fig. 3c). We also found similar increases in fold enrichment for marker subsets on chromosomes 21, 18, and 13, which are critical for NIPT detection (Supplementary Fig. 5).

We then expanded the analysis to a cohort of 28 plasma samples from pregnant women undergoing NIPT testing (median gestational age 10w6d, IQR 10w2d-11w3d). We sequenced these plasmas and blood cells more sparsely (median 0.8-1-fold coverage and median unique reads of 33M for HpaII and 46M for MspI), using only 0.5 mL of plasma (Supplementary Table 2). We divided the CpG enrichment markers into eight tiers based on their predicted enrichment performance. The 8 tiers demonstrated stepwise increases up to 5.6-fold in fetal fraction between the epiSIFT assay and the control (Fig. 3d, Supplementary Fig. 6a). With the most potent enrichment markers (tier T1), epiSIFT raised the median fetal fraction from 5.1% (IQR 4.3–6.3) before enrichment to 28.4% (IQR 24.4–33.0) after enrichment (Supplementary Table 3). Measured enrichment rose with model predictions across all 8 tiers, reaching 87% of the predicted gain at T1 (Fig. 3e, Supplementary Fig. 6b).

### Enrichment of Trisomy 21 in cfDNA Titrations

To test enrichment in an aneuploid background, we used the epiSIFT assay with enzymatic methylation conversion and computational enrichment of cfDNA titrations. We first created cfDNA Trisomy 21 titrations by physically titrating cfDNA (chorionic villus sampling culture media) from a fetus with Trisomy 21 (case BF4232) into plasma cfDNA from a healthy female donor (Fig. 4a). We performed epiSIFT and also performed methylation conversion to comprehensively read out methylation data across all CpG sites covered. Across measured titration levels ranging from 2% to 47%, the epiSIFT assay had an average 1.6-fold enrichment of fetal cfDNA (range: 1.3-1.9). Additional computational filtering based on the unmethylated status of the molecule increased enrichment to 3.0-fold (range: 2.0-4.1) (Fig. 4b). For example, an estimated low fetal fraction of 4% based on WGS increased to 15%, and a typical fetal fraction of 10% increased to 40%.

## Discussion

Achieving statistical significance in NIPT comes theoretically from two sources: fetal fraction and molecular counting. However, the latter is practically limited by dispersion, and the former has been difficult to increase genome-wide. Because fetal cfDNA is shorter than maternal cfDNA [15,20], size selection is one of the few established means of fetal enrichment. Selecting fragments below 160 bp roughly doubled fetal fraction (11.3% to 22.6%), cut test failures from 0.7% to 0.1%, and detected trisomies in 5 of 11 previously false-negative cases [21]. However, the two length distributions overlap substantially and overall DNA representation loss is significant, so the approach is not commonly implemented.

Methylation has previously served as a fetal marker, but with a limited number of hypermethylated foci rather than genome-wide purification [16,22]. Both approaches are constrained by a limited number of available hypermethylated placental markers [16]. Using high-depth whole methylome sequencing, our fetal methylome atlas has shown that hypomethylated markers vastly outnumber hypermethylated markers, to the point where we did not practically consider using hypermethylated markers (Fig. 2a–c).

Here we show that epigenetic enrichment is more discriminating between signal and noise and therefore more potent at enrichment than previous methods. Three components make that possible: (i) a nucleotide-resolution methylome of early fetal cfDNA, (ii) a model that ranks all CpGs as enrichment markers, and (iii) an assay that raised median fetal fraction from 5.1% to 28.4% in 28 plasma cases, and realized up to 13.7-fold enrichment in a single case at the most selective markers.

Discarded CVS culture media proved a surprisingly good source for the reference, yielding a median fetal purity of 94.7% at a depth of 207x across 26.3 million CpGs. Concordance with independently purified first-trimester trophoblast was high (ρ = 0.944) [9], and there is stability across karyotype, gestational age, fetal sex, and CVS samples.

The model uses fetal and background methylation at each CpG to rank markers by predicted enrichment potency. This ranking was supported in plasma, where fetal fraction increased across marker tiers in HpaII-enriched libraries but remained stable in MspI controls.

For aneuploidy screening, enrichment may be most valuable at low fetal fractions, where low fetal signal contributes to test failures and constrains lower-throughput platforms. By increasing fetal signal before sequencing, epiSIFT may reduce the depth needed for downstream analysis, though whether the gain translates into aneuploidy detection at reduced depth remains to be tested.

A few limitations should be considered. (i) Our fetal methylome reference was derived from cfDNA in CVS culture, where degradation limits its utility for fragmentomics or long read phasing. (ii) Our epiSIFT design addresses ~1.8 million CCGG motifs through the HpaII enzyme. While convenient, it is a minority of the total possible enrichment markers. Future additions of other methylation sensitive restriction enzymes may broaden this range. Enrichment selects a subset of molecules, increasing fetal fraction at the cost of molecule yield, and the degree of enrichment is uneven across the genome. Targeted sequencing could concentrate depth on the retained markers, although the optimal design will depend on the needs of the downstream application. (iii) The 28 plasma cohort was selected to quantify enrichment of fetal fraction rather than diagnostic performance of aneuploidy or other traits. For aneuploidy, we used Trisomy 21 titrations due to the lack of aneuploidy plasma cases.

Together, we establish a base resolution, first-trimester fetal cfDNA methylome, leverage it to create a precise enrichment model, and demonstrate physical enrichment of the fetal genome. Validation in native low-fetal-fraction and aneuploid plasma will determine the diagnostic value of epiSIFT.

## Methods

### Ethics statement

Residual discard materials including plasma and CVS culture supernatant were retrospectively included in this study under a waiver of informed consent approved by the Stanford Institutional Review Board (IRB #58461).

### Specimen collection

*In vitro* fetal cfDNA was recovered from the first culture media exchange after culturing a chorionic villus sample. Residual chorionic villus sampling culture media were collected at the point of discard from the Cytogenetics Laboratory at Stanford Health Care according to established clinical standard operating procedures. The samples were kept refrigerated at 4°C for a maximum of 5 days before processing. Upon receipt, samples were centrifuged at 16,000g for 10 minutes to pellet any residual cellular debris. The supernatant with cell-free DNA was carefully transferred using a pipette and stored in a −80°C freezer until DNA extraction.

### CVS or plasma fetal cfDNA

For cfDNA extraction, we used the Maxwell RSC ccfDNA Plasma Kit (AS1480 or AS1840 for large volume, Promega) to extract cfDNA from different sample types. All extractions were performed according to the manufacturer’s instructions, utilizing the Promega RSC 48 automated instrument. After extraction, cfDNA concentrations were quantified using the Qubit fluorometric assay (Q33231, Invitrogen). Purified DNA was then stored temporarily at 4°C or frozen at −20 to −80°C until downstream library preparation for sequencing.

### Sequencing CVS Methylome

We used EMseq or the NEBNext Enzymatic Methyl-seq Kit (E7120L, New England Biolabs) to create whole methylome libraries per the manufacturer’s instructions. Sequencing was conducted at the Broad Clinical Laboratories on the Illumina NovaSeqX platform using a 2×150 cycle format. For CVS cases BF4555 and upwards, we used 0.85x bead ratio rather than 1x ratio for the first bead cleaning to remove shorter molecules and generate longer sequencing libraries.

### EpiSIFT assay

FLEXseq was adapted to provide a physical implementation of epiSIFT. The FLEXseq method was previously described and detailed by Yu et al. [17] For epiSIFT, selection for unmethylated DNA was driven by replacing the MspI restriction enzyme in FLEXseq with the HpaII methylation-sensitive restriction enzyme that is highly selective for unmethylated CCGG motifs. The methylation conversion steps were optionally performed, and not performed for any experiment that involved SNP analysis.

As a brief overview of FLEXseq, we used components from the NEBNext Enzymatic Methyl-seq Kit (E7120L, New England Biolabs). Input DNA fragments were subjected to end repair and Adapter A ligation, followed by bead clean-up. DNA was next subjected to dephosphorylation, restriction enzyme digestion, end repair, and ligation with Adapter B. While all other steps and components are the same, we used 5 U HpaII (R0171M, New England Biolabs) instead of MspI. To allow for compatible buffers for the HpaII enzyme, rCutsmart buffer was used in place of Buffer 2 as described in the original FLEXseq protocol [17].

### Trisomy 21 cfDNA titration to demonstrate epiSIFT enrichment

Extracted fetal cfDNA (BF4232) was titrated into a healthy female donor (P143) extracted cfDNA. BF4232 was a first-trimester CVS specimen associated with a fetus with trisomy 21. Titrates were designed at fetal fractions of 30%, 10%, 3%, 1%, and 0.3%. Undiluted BF4232 cfDNA from CVS culture media and plasma cfDNA from the same donor were included as positive and negative references. Libraries created from the same titrated starting material were (i) unenriched whole-genome sequencing (WGS) libraries, (ii) epiSIFT libraries prepared with HpaII and (iii) FLEXseq control libraries prepared with MspI. Both epiSIFT and FLEXseq used enzymatic methylation conversion using the NEB Enzymatic Conversion components.

### Genome alignment and preprocessing

For the fetal cfDNA methylome, Enzymatic Methyl-seq (EM-seq) libraries were sequenced paired-end across multiple Illumina NovaSeq lanes. Sequencing adapters were trimmed with Cutadapt, and trimmed reads were aligned to the hg38 reference genome, as well as to the lambda and pUC19 control genomes, using Bismark v0.23.0. Reads exhibiting incomplete enzymatic conversion were removed by filtering out reads with unmethylated cytosines in the non-CpG context using the filter_non_conversion function (--consecutive --threshold 2). For each biological sample, alignments from all lanes were combined and PCR duplicates removed in a single pass with deduplicate_bismark (--multiple); samples prepared across two wells were deduplicated independently and the resulting deduplicated alignments were subsequently concatenated. Following non-conversion and duplicate filtering, methylation calls were extracted with the bismark_methylation_extractor function (--bedGraph), ignoring the first and last 10 bp of both reads (--ignore 10 --ignore_r2 10 --ignore_3prime 10 --ignore_3prime_r2 10) to remove end-repair–associated methylation bias characteristic of EM-seq, following best practices for processing EM-seq data. The methylation calls of each CpG dinucleotide from the top and bottom strands, staggered by 1 bp, were merged using the coverage2cytosine function (--merge_CpG --discordance 50 --zero_based). Finally, common single-nucleotide polymorphism positions (dbSNP153Common.bed dataset from dbSNP Build 153 with frequency ≥ 1% in the 1000 Genomes Project) were removed using BEDTools to filter out artifacts and for sample de-identification.

For WGBS background plasma data, we downloaded the hg38 BETA files directly from the Loyfer et al.[11] WGBS data (GSE186458) and converted them to a BED format.

For each FLEXseq (MspI) or epiSIFT (HpaII) sample with methylation conversion, read 1 (R1) and read 2 (R2) files from different sequencing lanes were merged separately prior to downstream processing. Adapter trimming and quality filtering were performed using Cutadapt, with the 3′ adapter sequences for R1 and R2 specified as NNAGATCGGAAGAGC. High-quality reads were subsequently aligned to the human reference genome (GRCh38/hg38) using Bismark (v0.23.0) with default settings. To ensure accurate methylation calling, reads exhibiting incomplete methylation conversion—defined as containing more than three methylated cytosines in a non-CG context—were excluded using the filter_non_conversion function in Bismark. Duplicate reads, including those potentially arising from PCR amplification, were identified and removed using the deduplicate_bismark script. The resulting BAM files, comprising non-redundant and methylation-converted reads, were used for downstream methylation and genomic analyses.

Alignment and processing of each FLEXseq (MspI) or epiSIFT (HpaII) sample without methylation conversion involve first adapter trimming described above. FASTQ reads were then aligned with bwa mem (v0.7.17) with the settings -K 100000000 -Y, ALT-aware to GRCh38/hg38. Duplicate marking and base quality score recalibration were done by GATK 4.6.2.0.

### Integrated CVS methylome reference and validation

CpG-level methylation coverage files generated from CVS culture-media EM-seq libraries were used to construct an integrated fetal cfDNA methylome reference. Sample inclusion for reference construction and validation analyses was controlled using analysis-specific sample lists. For each CpG, methylated and unmethylated read counts were summed across the 28 CVS samples included in the reference, and the integrated beta value was calculated as methylated reads divided by total reads. Analyses were restricted to standard hg38 autosomes and sex chromosomes. The per-sample CVS CpG coverage files used for reference construction had been filtered to remove CpGs overlapping common SNP positions.

Fetal cfDNA purity was estimated from whole-genome sequencing copy-number profiles generated with CNVkit. For CVS media samples with fetal trisomy or male fetal sex, segment-level log2 copy ratios from the relevant chromosome were used to estimate the fraction of fetal/placental DNA in the sample. For trisomies, purity was calculated as (1 - 2^(log2 ratio)) / (1 - 3/2). For chromosome X copy number in male fetuses or the monosomy-X sample BF4388, purity was calculated as (1 - 2^(log2 ratio)) / (1 - 1/2). When both trisomy-based and chromosome-X-based estimates were available, the two estimates were averaged.

Fragment-length distributions were derived from whole-genome sequencing data by summarizing paired-end insert sizes from deduplicated alignments. The R33 and R34 WGS workflows used different bead-cleanup conditions; the R34 cleanup selected against shorter fragments. Samples were therefore stratified as R33 (n=17) or R34 (n=11), and the two rounds were not pooled for fragment-length interpretation. Within each round, per-sample insert-size distributions were converted to percent frequencies and averaged so that each sample contributed equally. The modal fragment length was defined as the insert size at the maximum of the resulting cohort frequency curve; curves were smoothed with a Gaussian kernel (σ = 2 bp) for visualization.

Internal consistency of the CVS methylome reference was evaluated by comparing group-level methylation profiles across fetal karyotype and gestational age, as well as fetal sex. For the trisomy 21 comparison, 13 T21 and 11 normal karyotype samples were aggregated into separate groups. For the gestational-age comparison, the 28 CVS samples included in the reference were divided at the median gestational age into earlier- and later-gestation groups of 14 samples each. The fetal-sex comparison included 12 XX and 14 XY samples, as two XXY samples were excluded from this binary comparison. In all three analyses, CpG-level methylated and unmethylated read counts were summed within each group, group-level beta values were calculated, and CpGs were retained if both groups had minimum aggregated depth of 10. Each sample was also compared with a leave-one-out reference constructed from the remaining 27 samples; correlations used CpGs with depth of at least 5 in the held-out sample and at least 50 in the leave-one-out reference. Pearson correlations were calculated across all eligible CpGs, whereas Spearman’s rho was calculated using a reproducible random subset of 1,000,000 eligible CpGs for each sample.

The integrated CVS reference was compared with purified first-trimester trophoblast methylation profiles from Yuan et al. Probe-level beta values from the five available first-trimester purified trophoblast samples were averaged to create a Yuan trophoblast reference. Probes were mapped to hg38 genomic coordinates using the EPIC hg38 manifest and overlapped with the integrated CVS reference at CpGs with integrated CVS depth of at least 10. Pearson and Spearman correlations were calculated from paired CpG-level beta values before graphical binning.

All reference construction, sample selection, methylation comparisons, correlation analyses, and figure-generation steps were implemented as reproducible Python scripts to be released with the paper. Scripts were run in a documented Python environment using pandas, NumPy, SciPy, and Matplotlib. Density heatmaps were generated by binning paired methylation percentages across the 0-100% range. Figure 1 density heatmaps used 100 × 100 bins and a logarithmic CpG-count color scale capped at 10^4 CpGs per bin.

### Calculating and analyzing the enrichment model

The integrated 28-sample CVS methylome reference was compared with a normal plasma cfDNA methylome reference generated from Loyfer et al. per-sample normal plasma methylation BED files. For each reference, methylated and total read counts were summed at each CpG, and beta values were calculated as methylated reads divided by total reads. CpGs were matched by exact hg38 chromosome, start, and end coordinate. CVS-vs-normal analyses retained CpGs with integrated CVS depth of at least 50 and integrated normal plasma depth of at least 450.

Density heatmaps were generated from paired beta values after converting them to percent methylation for display, with axes spanning 0-100% methylation. Pearson and Spearman correlations were calculated from paired CpG-level beta values before graphical binning. Density heatmaps used logarithmic CpG-count color scales.

Theoretical fetal enrichment was modeled for each paired CpG at a fetal fraction of 4%, using the integrated CVS reference as the fetal/placental methylation profile and the Loyfer et al. normal plasma reference as the background plasma methylation profile. Hypomethylated fetal markers were defined as CpGs with normal plasma beta greater than CVS beta. For these sites, theoretical fold enrichment was calculated as (1 - CVS beta) / (((1 - CVS beta) × f) + ((1 - normal plasma beta) × (1 - f))), where f = 0.04. For hypermethylated fetal markers, defined as CpGs with CVS beta greater than normal plasma beta, fold enrichment was calculated as CVS beta / ((CVS beta × f) + (normal plasma beta × (1 - f))). Fold-enrichment contours were overlaid only in the hypomethylated-marker region. The 1,839,680 CCGG sites in the reference were ranked genome-wide by predicted fold enrichment and separated into octiles of ~229,960 CpGs each, designated T1 (highest) to T8 (lowest).

Marker-yield curves were generated by counting hypomethylated CpGs with theoretical fold enrichment greater than or equal to each threshold from 0 to 8, separately for the whole genome and chromosome X. Uncapped fold-enrichment values were used for these threshold calculations.

For genomic-context analysis, non-neutral CpGs were ranked within each chromosome by direction-specific fold enrichment, and the top 50%, 5%, and 0.1% were selected, rounding up, before separation by methylation direction. CpGs overlapping hg38 GENCODE v36 annotations by at least 1 bp were assigned mutually exclusively, in priority order, to MANE Select promoter (−1.5/+0.5 kb from the transcription start site), coding sequence, UTR, other exon, intron, or intergenic; percentages were calculated within each direction and subset. The same assignment was applied to the T1–T8 octiles (Supplementary Fig. 4a). Enhancer overlap used FANTOM5 reprocessed hg38 permissive enhancers [23] (63,280 merged intervals) under the same ≥1 bp rule, scored independently (Supplementary Fig. 4b).

Chromosome-level fold-enrichment distributions were plotted for hypomethylated CpGs passing the same CVS and normal plasma depth filters. Fold-enrichment values were grouped by chromosome and displayed as violin plots with internal boxplots; values above 10 were capped for visualization only.

Script generation and pipeline development were aided by ChatGPT Codex 5.6 Sol model, and manually verified.

### Simulated Enrichment Model and Equation

In this model, we assumed that each DNA molecule could be assigned as target or background based on its methylation status. For each CpG biomarker we defined the methylation fraction (or beta) in the target (CVS; β_T_) and background (normal plasma; β_B_) as methylated reads divided by total reads. We then compared the two: where the target was less methylated than background (β_T_ < β_B_), the biomarker is considered hypomethylated and unmethylated reads are assigned to the target DNA group. Conversely, if the target was more methylated than the background (β_T_ > β_B_), the biomarker is considered hypermethylated and methylated reads are selected. We model the sample as a mixture in which a fraction p of molecules is target and 1 − p is background (1 represents 100% of molecules). A read is retained if it matches the selected state, so the target is retained at rate (1 − β_T_) for hypomethylated sites (or β_T_ for hypermethylated), and background at rate (1 − β_B_) (or β_B_ for hypermethylated). Fold enrichment is the target retention rate divided by the average retention rate of the whole sample, giving Equation 1 for hypomethylated sites and Equation 2 for hypermethylated sites.

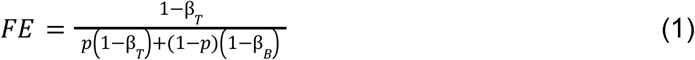

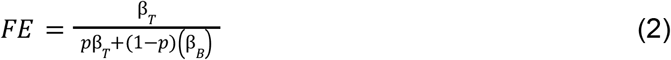

Setting p = 0 recovers the maximum attainable enrichment, (1 − β_T_)/(1 − β_B_) for hypomethylated sites, while larger p lowers fold enrichment (FE) as target molecules increasingly dilute their own selection. Reported values assume a 4% fetal target fraction or p = 0.04. FE > 1 indicates enrichment, FE = 1 none, and FE < 1 depletion.

### Analysis: Determining the maternal genotypes

After alignment, maternal cell libraries prepped with FLEXseq (MspI without conversion) were called with bcftools mpileup with settings -m -C alleles (bcftools 1.19). A minimum maternal depth of 10 was used, with measured depth in these libraries approximately 6. ChrX was excluded because male fetuses inherit chromosome X only from the mother and would therefore not have informative paternal alleles. Sites of interest were recognized SNP positions (population frequency >0.1 from gnomAD r2.0.1), where the maternal state was homozygous, because an alternate allele then counts as fetal, coming from a paternal origin. SNPs that create or destroy a CCGG site were excluded as they would be a confounder.

### Analysis: Estimating fetal fraction from single-nucleotide polymorphisms (SNP)

FLEXseq (MspI) and epiSIFT (HpaII) libraries without methylation conversion were inputs. Alignment to hg38 using BWA yielded BAM files, which were deduplicated. When using a subset of CpG markers, a BED file is first created as a mask for the markers of interest from the Enrichment model (all markers, a top-ranked fraction, or a single tier). Next, a custom Python script filters the deduplicated BAM file with this mask to only retain reads whose CCGG selector CpGs overlapped with the mask. Plasma allele counts were tallied using samtools mpileup (samtools 1.3.1) with -q 30 -Q 20 -d 1000. Overlapping paired-end mates were adjusted so that overlaps were not double counted.

To estimate the fetal fraction, a binomial-mixture maximum likelihood estimator (MLE) was used as previously described[18,19]. The MLE inputs were population frequencies based on gnomAD frequencies, maternal homozygous SNP positions as determined by the detected maternal genotypes from the maternal cells and their allelic counts in plasma, and a global error rate. Assuming a relatively constant error rate, the error rate is first calculated across SNPs using alternative bases not identified in the population alleles as previously described[19]. Fold-enrichment was then calculated by dividing the fetal fraction of epiSIFT (using HpaII enzyme) by FLEXseq (using MspI enzyme). No depth cut-off was used; instead all available SNP positions were integrated for the estimate.

For each tier, we converted the median CpG-level prediction from its assumed 4% fetal fraction to each case’s SNP-based MspI fetal fraction estimate using the enrichment model. Fig. 3e uses cohort median MspI fraction for prediction and the ratio of cohort median fetal fractions for measurement. Gain recovery was calculated as (measured enrichment − 1)/(predicted enrichment − 1).

Script generation was aided by Claude Code Opus 5 model and manually verified.

### Analysis: Trisomy 21 cfDNA titration

We described the titration under the “Trisomy 21 cfDNA titration” section above. To analyze the titration enrichment by epiSIFT, we sourced two samples: normal adult female plasma in the 25-35 age range (P143) to represent the pure maternal plasma, and a chorionic villus sample (BF4232) with trisomy-21 to help determine the exact fetal purity (~91%) and titration. Unenriched control sequencing library preparation used MspI (FLEXseq), which cleaves all C^CGG regardless of methylation status. The physically enriched (epiSIFT) library preparation used HpaII, which cleaves C^CGG only once when the CpG is unmethylated.

We estimated the fetal fraction for each titration from WGS libraries using chromosome 21 copy number. Chromosome 21 log_2_ ratios were obtained from GC- and mappability-corrected read depth in 500-kb bins (ichorCNA), and each library was summarized as the median log_2_ ratio r across the 56 bins spanning chromosome 21q (14.0-46.5 Mb); the short arm was excluded owing to negligible reference depth. The fraction of trisomic DNA was calculated as (1 − 2^r)/(1 − 3/2), the same equation used for trisomy-based CVS fetal purity. This gives the fraction of trisomy placental DNA rather than the fraction of culture-media cfDNA added, so it already incorporates the 91% purity of the undiluted library, measured by the same equation. The WGS fractions were 47%, 24%, 10%, 4%, and 2% for the planned 30%, 10%, 3%, 1%, and 0.3% fractions respectively, presumably due to differences in the lengths of the starting material.

For computational epiSIFT filtering, read pairs from each HpaII library were restricted to the hypomethylation marker panel (result_methy_panelA.bed, 12,542,345 CpGs) with bedtools pairtobed, and only fully unmethylated fragments were retained using Bismark methylation_consistency (--min-count 1 --paired_end). Fetal fraction was estimated for the epiSIFT libraries before and after this filtering by the chromosome 21 method above, using a panel of normals built from the matched unenriched libraries. Fold enrichment at each titration level is the ratio of the epiSIFT fetal fraction to the WGS fetal fraction for that level.

## Code and Data Availability

Data and analysis scripts will be deposited in appropriate repositories upon publication. Public references used include GSE186458 (Loyfer et al.) and GSE159526 (Yuan et al.).

