## Supplemental Tables 1-3 for "Early Prenatal Cell-Free DNA Methylation Landscape Informs Epigenetic Purification of the Circulating Fetal Genome"

### Supplementary Table 1. Chorionic villus sampling (CVS) case characteristics and sequencing metrics

*Per-case demographic, sequencing, purity, and methylation conversion quality-control metrics for the 30 CVS specimens sequenced in this study. Twenty-nine were processed through the EM-seq reference-building pipeline, of which 28 were used to build the integrated fetal cfDNA methylome reference (Fig. 1). Fetal purity could be estimated in the 23 cases with an informative trisomy or sex-chromosome copy-number change. The two specimens not included in the reference are shaded: BF4388, excluded for low fetal purity, and BF4232, which was not processed through the reference-building pipeline and served as the source of trisomy 21 cfDNA for the titration experiment (Fig. 4).*

| Case ID (BF) | Maternal Age (range) | Gestational Age | Fetal Sex | Fetal Genetics or Karyotype | Total Sequencing Reads | Total Deduplicated Reads | Fetal Purity (%)[d] | Methylation False Positive Rate (%)[a] | Methylation False Negative Rate (%)[b] | Reference Cohort Status[c] |
|---|---|---|---|---|---|---|---|---|---|---|
| *BF4232* | *35-39* | *13w3d* | *XY* | *Trisomy 21* | *–* | *–* | *90.7* | *–* | *–* | *Not included, used for titration* |
| BF4371 | 40-44 | 12w4d | XY | Trisomy 21 | 131,309,394 | 101,559,548 | 98.4 | 0.2 | 2.8 | Included |
| *BF4388* | *30-34* | *12w2d* | *X0* | *Monosomy X* | *218,004,860* | *100,778,928* | *18.7* | *0.15* | *3.0* | *Excluded for low purity* |
| BF4479 | 25-29 | 14w4d | XY | Normal | 158,730,774 | 120,306,982 | 100.0 | 0.2 | 2.4 | Included |
| BF4500 | 30-34 | 13w4d | XY | Normal | 144,212,258 | 112,158,074 | 90.3 | 0.1 | 3.8 | Included |
| BF4501 | 20-24 | 13w | XY | Normal | 135,962,414 | 103,714,410 | 105.6 | 0.2 | 3.6 | Included |
| BF4502 | 40-44 | 13w | XY | Trisomy 21 | 271,640,690 | 210,457,204 | 98.7 | 0.15 | 3.2 | Included |
| BF4511 | 35-39 | 13w | XY | Normal | 287,492,952 | 220,707,548 | 86.8 | 0.2 | 1.9 | Included |
| BF4512 | 40-44 | 12w2d | XX | Trisomy 21 | 235,316,750 | 183,147,774 | 92.5 | 0.2 | 2.3 | Included |
| BF4530 | 35-39 | 13w | XX | Normal | 147,284,038 | 115,812,056 | Not Measured | 0.2 | 2.3 | Included |
| BF4531 | 40-44 | 12w6d | XY | Trisomy 18 | 293,755,004 | 223,946,644 | 101.9 | 0.2 | 1.9 | Included |
| BF4532 | 30-34 | 12w2d | XXY | XXY | 130,920,174 | 101,652,860 | Not Measured | 0.1 | 3.6 | Included |
| BF4533 | 35-39 | 13w6d | XX | Trisomy 21 | 318,525,566 | 246,748,122 | 92.8 | 0.25 | 2.1 | Included |
| BF4534 | 40-44 | 12w3d | XY | Normal | 134,109,818 | 103,286,818 | 97.1 | 0.2 | 4.0 | Included |
| BF4541 | 35-39 | 13w | XX | Trisomy 18 | 147,576,096 | 113,844,382 | 93.8 | 0.4 | 1.0 | Included |
| BF4542 | 40-44 | 13w | XX | Trisomy 21 | 304,744,996 | 238,069,790 | 90.8 | 0.25 | 2.1 | Included |
| BF4543 | 35-39 | 13w6d | XXY | XXY | 184,408,036 | 143,653,820 | Not Measured | 0.3 | 2.0 | Included |
| BF4548 | 30-34 | 13w6d | XX | Trisomy 21 | 134,811,478 | 107,109,962 | 92.4 | 0.2 | 2.7 | Included |
| BF4549 | 35-39 | 12w5d | XY | Trisomy 21 | 283,413,828 | 218,966,190 | 99.0 | 0.25 | 2.7 | Included |
| BF4555 | 35-39 | 12w5d | XX | Trisomy 21 | 150,931,642 | 116,124,014 | 94.4 | 0.2 | 2.7 | Included |
| BF4556 | 35-39 | 12w | XX | Trisomy 21 | 151,117,256 | 116,694,668 | 94.7 | 0.2 | 3.2 | Included |
| BF4557 | 35-39 | 11w6d | XY | Trisomy 21 | 136,801,826 | 105,211,554 | 94.5 | 0.3 | 2.1 | Included |
| BF4564 | 35-39 | 12w0d | XX | Normal | 154,669,188 | 118,908,648 | Not Measured | 0.2 | 2.6 | Included |
| BF4570 | 35-39 | 12w3d | XY | Trisomy 21 | 156,132,696 | 119,078,374 | 99.2 | 0.2 | 3.4 | Included |
| BF4579 | 25-29 | 12w4d | XX | Trisomy 21 | 118,307,668 | 93,294,282 | 90.4 | 0.3 | 1.8 | Included |
| BF4580 | 35-39 | 11w2d | XX | Normal | 132,796,616 | 104,027,200 | Not Measured | 0.1 | 1.9 | Included |
| BF4581 | 35-39 | 11w2d | XY | Trisomy 21 | 145,471,594 | 112,367,766 | 96.3 | 0.2 | 3.6 | Included |
| BF4585 | 35-39 | 11w4d | XX | Normal | 142,300,590 | 109,778,198 | Not Measured | 0.2 | 3.0 | Included |

### Supplementary Table 1. Chorionic villus sampling (CVS) case characteristics and sequencing metrics

*Per-case demographic, sequencing, purity, and methylation conversion quality-control metrics for the 30 CVS specimens sequenced in this study. Twenty-nine were processed through the EM-seq reference-building pipeline, of which 28 were used to build the integrated fetal cfDNA methylome reference (Fig. 1). Fetal purity could be estimated in the 23 cases with an informative trisomy or sex-chromosome copy-number change. The two specimens not included in the reference are shaded: BF4388, excluded for low fetal purity, and BF4232, which was not processed through the reference-building pipeline and served as the source of trisomy 21 cfDNA for the titration experiment (Fig. 4).*

| Case ID (BF) | Maternal Age (range) | Gestational Age | Fetal Sex | Fetal Genetics or Karyotype | Total Sequencing Reads | Total Deduplicated Reads | Fetal Purity (%)[d] | Methylation False Positive Rate (%)[a] | Methylation False Negative Rate (%)[b] | Reference Cohort Status[c] |
|---|---|---|---|---|---|---|---|---|---|---|
| BF4586 | 30-34 | 13w0d | XY | Normal | 125,270,160 | 98,022,352 | 99.3 | 0.2 | 4.8 | Included |
| BF4592 | 30-34 | 13w0d | XY | Normal | 148,485,068 | 114,585,214 | 98.2 | 0.3 | 3.1 | Included |

**Notes**

a. Methylation False Positive Rate = % of cytosines in the unmethylated Lambda spike-in control called as methylated (CpG context), reflecting incomplete enzymatic conversion. Both rates are calculated after non-conversion filtering.

b. Methylation False Negative Rate = % of cytosines in the CpG-methylated pUC19 spike-in control called as unmethylated (CpG context), reflecting over-conversion.

c. Reference cohort status. BF4388 (18.7% estimated fetal purity) was processed through the EM-seq reference-building pipeline but excluded from the integrated CVS methylome reference due to low fetal purity. Its sequencing and quality-control metrics are reported here for completeness. BF4232 was not processed through the reference-building pipeline and was used exclusively as the trisomy 21 donor for the cfDNA titration experiment (Fig. 4). Dashes indicate metrics that are not applicable because no EM-seq methylome library was prepared for this specimen.

d. Values slightly exceeding 100% reflect estimator variance at shallow depth. "Not Measured" indicates cases with no informative trisomy or sex-chromosome copy-number change. See Methods for the estimation procedure. For BF4232, fetal purity was estimated from chromosome 21 copy number in the undiluted culture-media library as part of the titration workflow rather than the workflow applied to the reference-cohort specimens.

### Supplementary Table 2. Maternal plasma case characteristics and sequencing quality-control metrics

*Per-case demographics, sequencing depth, and quality-control metrics for the 28-case maternal plasma cohort used in the epiSIFT enrichment experiments (Fig. 3d,e). Two additional plasma samples, shaded, are not part of that cohort: P352, a deep-sequenced single case obtained through beta-human chorionic gonadotropin testing in the Emergency Department rather than the NIPT collection pathway (Fig. 3a–c), and P143, a non-pregnant female plasma donor used as the background for the cfDNA titration experiment (Fig. 4).*

| Case ID | Fetal Sex[a] | Maternal Age (range) | Gestational Age | Total Reads HpaII | Total Reads MspI | Total Deduplicated Reads HpaII | Total Deduplicated Reads MspI | On-target CCGG % HpaII[b] | On-target CCGG % MspI[b] | Plasma Cohort Status[c] |
|---|---|---|---|---|---|---|---|---|---|---|
| *P143* | *–* | *25-29* | *–* | *–* | *–* | *31,795,497* | *97,869,566* | *93.0* | *98.8* | *Not included, plasma titration control* |
| *P352* | *–* | *40-44* | *–* | *1,588,345,052* | *756,393,474* | *62,754,658* | *115,090,596* | *91.2* | *97.9* | *Not included, single-case analysis* |
| P673 | M | 30-34 | 10w2d | 50,726,320 | 60,781,774 | 33,302,581 | 46,130,135 | 90.7 | 97.2 | Included |
| P679 | M | 35-39 | 10w6d | 53,235,870 | 69,986,580 | 34,606,220 | 51,189,738 | 92.5 | 97.3 | Included |
| P683 | F | 30-34 | 11w0d | 54,888,314 | 76,913,986 | 34,250,633 | 55,519,943 | 92.3 | 97.4 | Included |
| P686 | M | 30-34 | 11w3d | 52,125,160 | 64,894,988 | 31,474,259 | 46,345,104 | 92.7 | 97.1 | Included |
| P689 | M | 35-39 | 10w6d | 46,335,592 | 58,750,606 | 20,552,880 | 34,407,752 | 91.1 | 96.9 | Included |
| P690 | M | 35-39 | 10w0d | 55,588,006 | 72,079,000 | 36,671,778 | 54,565,506 | 92.4 | 97.2 | Included |
| P693 | M | 40-44 | 11w5d | 55,303,482 | 55,746,404 | 32,089,979 | 40,026,614 | 92.2 | 97.2 | Included |
| P695 | M | 20-24 | 11w6d | 55,775,156 | 76,198,342 | 33,613,491 | 53,062,935 | 91.9 | 97.3 | Included |
| P705 | M | 25-29 | 10w3d | 64,317,630 | 68,177,942 | 36,670,420 | 48,233,904 | 93.4 | 97.3 | Included |
| P706 | M | 25-29 | 11w4d | 62,937,700 | 73,476,526 | 29,352,968 | 42,203,217 | 95.0 | 97.1 | Included |
| P709 | M | 30-34 | 10w2d | 57,432,172 | 72,190,868 | 29,927,885 | 46,743,072 | 93.8 | 97.2 | Included |
| P710 | M | 35-39 | 10w2d | 56,434,184 | 76,885,610 | 32,337,406 | 52,454,421 | 93.8 | 97.3 | Included |
| P711 | M | 35-39 | 11w3d | 47,539,276 | 59,373,432 | 27,963,724 | 41,466,368 | 93.2 | 97.3 | Included |
| P719 | M | 30-34 | 13w5d | 48,963,350 | 62,076,124 | 29,098,909 | 44,260,676 | 93.4 | 97.1 | Included |
| P721 | M | 25-29 | 10w0d | 77,339,162 | 55,441,178 | 43,705,496 | 41,112,134 | 93.0 | 97.1 | Included |
| P722 | M | 40-44 | 10w1d | 52,658,810 | 54,869,246 | 34,859,017 | 43,175,166 | 91.5 | 97.3 | Included |
| P727 | M | 30-34 | 10w4d | 58,167,236 | 52,458,284 | 38,214,671 | 41,170,235 | 90.9 | 97.3 | Included |
| P731 | M | 30-34 | 10w1d | 57,892,082 | 69,242,572 | 39,753,279 | 53,137,511 | 91.5 | 97.1 | Included |
| P735 | M | 40-44 | 10w6d | 57,190,006 | 67,944,468 | 33,843,569 | 47,435,281 | 93.1 | 97.1 | Included |
| P736 | M | 35-39 | 10w0d | 50,008,762 | 87,415,340 | 33,302,429 | 62,648,117 | 91.8 | 97.2 | Included |
| P765 | M | 35-39 | 11w0d | 46,275,548 | 57,809,162 | 20,307,650 | 34,564,185 | 94.4 | 96.8 | Included |

### Supplementary Table 2. Maternal plasma case characteristics and sequencing quality-control metrics

*Per-case demographics, sequencing depth, and quality-control metrics for the 28-case maternal plasma cohort used in the epiSIFT enrichment experiments (Fig. 3d,e). Two additional plasma samples, shaded, are not part of that cohort: P352, a deep-sequenced single case obtained through beta-human chorionic gonadotropin testing in the Emergency Department rather than the NIPT collection pathway (Fig. 3a–c), and P143, a non-pregnant female plasma donor used as the background for the cfDNA titration experiment (Fig. 4).*

| Case ID | Fetal Sex[a] | Maternal Age (range) | Gestational Age | Total Reads HpaII | Total Reads MspI | Total Deduplicated Reads HpaII | Total Deduplicated Reads MspI | On-target CCGG % HpaII[b] | On-target CCGG % MspI[b] | Plasma Cohort Status[c] |
|---|---|---|---|---|---|---|---|---|---|---|
| P770 | M | 30-34 | 10w5d | 54,054,932 | 64,988,062 | 33,421,364 | 46,933,461 | 92.6 | 97.2 | Included |
| P778 | M | 30-34 | 10w6d | 52,975,450 | 69,693,566 | 27,834,222 | 43,566,448 | 93.0 | 96.7 | Included |
| P781 | M | 30-34 | 12w4d | 56,736,148 | 64,004,970 | 31,875,023 | 44,263,037 | 93.1 | 97.4 | Included |
| P782 | M | 30-34 | 10w2d | 56,526,638 | 63,951,202 | 22,503,675 | 34,980,573 | 95.7 | 97.0 | Included |
| P786 | M | 30-34 | 11w3d | 64,170,368 | 82,421,258 | 33,195,097 | 51,283,694 | 93.8 | 97.2 | Included |
| P796 | M | 30-34 | 12w0d | 52,640,062 | 74,333,660 | 33,819,091 | 52,668,265 | 92.5 | 97.1 | Included |
| P805 | M | 30-34 | 11w0d | 48,094,828 | 61,596,826 | 29,257,065 | 43,455,339 | 93.9 | 97.2 | Included |

**Notes**

a. Fetal sex was determined from clinical NIPT (Natera) results where available, or from the delivery record once the pregnancy reached delivery for cases where NIPT sex reporting was not available.

b. On-target CCGG % = percentage of reads with a CCGG motif at the read 1 5′ end (±2 bp), assessed on chromosome 1, reflecting enzymatic cutting specificity of the FLEXseq/epiSIFT library preparation.

c. Cohort status. The 28 cases marked "Included" comprise the maternal plasma cohort (Fig. 3d,e). P352, obtained outside the NIPT collection pathway, is reported individually in the main text and figures. P143 is a non-pregnant female donor, so the age shown is donor age. Dashes indicate values that are not available or not applicable.

### Supplementary Table 3. Fetal fraction by CpG enrichment marker tier in the maternal plasma cohort, control versus epiSIFT-enriched

*Per-case fetal fraction for all 28 plasma cohort cases in the unenriched control channel (MspI) and the epiSIFT-enriched channel (HpaII), for the unfiltered library and for each of eight CpG marker tiers ordered from lowest (T8) to highest (T1) predicted fold enrichment. Clinical NIPT fetal fraction (Natera) is shown for comparison where reported (n=19), but not involved in enrichment calculations. Source data for Fig. 3d,e and Supplementary Fig. 6.*

| Case characteristics | | | | Control (MspI)[a,c] | | | | | | | | | epiSIFT-enriched (HpaII)[a,c,d] | | | | | | | | |
|---|---|---|---|---|---|---|---|---|---|---|---|---|---|---|---|---|---|---|---|---|---|
| Case ID | Fetal Sex | Gestational Age | Natera Fetal Fraction[e] | Unfiltered[b] | T8 | T7 | T6 | T5 | T4 | T3 | T2 | T1 | Unfiltered[b] | T8 | T7 | T6 | T5 | T4 | T3 | T2 | T1 |
| P683 | F | 11w0d | Not available | 4.6% | 4.4% | 4.5% | 4.3% | 4.0% | 3.7% | 3.8% | 4.0% | 4.1% | 7.8% | 3.6% | 5.9% | 8.3% | 10.8% | 12.2% | 15.2% | 19.9% | 27.6% |
| P711 | M | 11w3d | Not available | 4.7% | 4.3% | 4.3% | 4.5% | 4.1% | 3.5% | 3.9% | 4.2% | 3.5% | 6.6% | 3.7% | 4.9% | 6.1% | 8.3% | 10.1% | 11.3% | 14.1% | 21.6% |
| P781 | M | 12w4d | 4.5% | 4.7% | 4.2% | 4.3% | 4.5% | 4.0% | 3.7% | 3.6% | 3.7% | 3.9% | 8.2% | 4.2% | 5.4% | 8.4% | 9.9% | 12.2% | 15.7% | 22.2% | 28.9% |
| P709 | M | 10w2d | Not available | 4.7% | 4.6% | 4.4% | 4.1% | 4.1% | 3.5% | 4.2% | 4.0% | 3.6% | 7.3% | 3.8% | 5.8% | 7.2% | 9.8% | 12.4% | 13.8% | 15.0% | 19.4% |
| P805 | M | 11w0d | 6.7% | 4.8% | 4.1% | 4.4% | 4.4% | 4.2% | 4.3% | 4.3% | 4.0% | 4.3% | 7.4% | 3.9% | 5.3% | 7.2% | 10.7% | 13.4% | 13.3% | 17.7% | 22.6% |
| P689 | M | 10w6d | 4.8% | 4.9% | 4.2% | 4.3% | 4.1% | 4.1% | 3.7% | 3.9% | 4.2% | 4.6% | 6.2% | 3.9% | 4.6% | 5.6% | 8.3% | 8.3% | 10.9% | 11.9% | 21.9% |
| P705 | M | 10w3d | 8.0% | 4.9% | 4.5% | 4.5% | 4.2% | 4.1% | 4.1% | 4.1% | 4.3% | 4.5% | 9.2% | 4.3% | 5.9% | 9.0% | 13.4% | 15.1% | 17.0% | 21.3% | 26.7% |
| P782 | M | 10w2d | 8.7% | 4.9% | 5.2% | 5.4% | 4.7% | 4.2% | 3.0% | 3.3% | 3.1% | 2.9% | 5.7% | 4.4% | 5.1% | 4.5% | 5.2% | 6.9% | 7.6% | 12.8% | 13.2% |
| P786 | M | 11w3d | Not available | 5.0% | 4.1% | 5.2% | 4.7% | 4.8% | 4.3% | 4.5% | 4.7% | 4.5% | 7.1% | 3.8% | 5.7% | 6.2% | 10.2% | 10.5% | 14.5% | 17.2% | 22.7% |
| P721 | M | 10w0d | 10.4% | 5.1% | 4.3% | 5.1% | 5.1% | 4.8% | 4.5% | 4.2% | 5.0% | 4.3% | 10.4% | 4.7% | 6.9% | 9.9% | 13.6% | 15.5% | 19.7% | 25.2% | 27.1% |
| P765 | M | 11w0d | 9.2% | 5.2% | 5.0% | 4.9% | 4.5% | 4.6% | 4.2% | 4.3% | 4.4% | 4.1% | 7.1% | 4.1% | 5.4% | 8.0% | 9.5% | 11.7% | 15.2% | 16.4% | 32.6% |
| P690 | M | 10w0d | 7.0% | 5.2% | 4.5% | 4.9% | 4.5% | 4.8% | 4.4% | 4.1% | 5.0% | 4.7% | 10.0% | 4.3% | 6.8% | 9.0% | 13.4% | 15.4% | 16.0% | 22.3% | 25.0% |
| P736 | M | 10w0d | 6.1% | 5.3% | 4.7% | 5.2% | 5.0% | 5.2% | 4.3% | 4.2% | 4.4% | 4.5% | 8.0% | 3.6% | 5.6% | 8.9% | 11.2% | 13.9% | 13.4% | 19.5% | 22.8% |
| P719 | M | 13w5d | 12.2% | 5.6% | 5.1% | 5.7% | 5.3% | 4.8% | 5.0% | 4.8% | 5.4% | 5.2% | 9.7% | 4.6% | 7.5% | 10.5% | 12.8% | 15.7% | 17.1% | 24.7% | 25.2% |
| P727 | M | 10w4d | Not available | 5.7% | 5.3% | 5.8% | 6.1% | 5.4% | 4.8% | 5.3% | 5.4% | 4.9% | 10.8% | 4.6% | 7.2% | 11.2% | 16.6% | 17.4% | 20.3% | 24.1% | 34.9% |
| P735 | M | 10w6d | 11.0% | 6.1% | 5.7% | 6.2% | 5.9% | 5.3% | 5.4% | 5.4% | 5.6% | 5.3% | 10.7% | 5.5% | 7.9% | 11.2% | 14.3% | 16.4% | 17.9% | 24.5% | 29.3% |
| P710 | M | 10w2d | 10.4% | 6.2% | 5.0% | 6.0% | 6.1% | 5.1% | 5.3% | 5.8% | 5.5% | 5.2% | 10.1% | 4.9% | 7.4% | 10.7% | 13.4% | 17.2% | 19.5% | 23.9% | 28.0% |
| P706 | M | 11w4d | 12.9% | 6.3% | 5.6% | 5.9% | 6.0% | 5.2% | 5.5% | 5.5% | 5.9% | 6.0% | 10.4% | 5.3% | 7.7% | 10.9% | 15.7% | 14.9% | 22.1% | 21.4% | 34.1% |
| P722 | M | 10w1d | 6.5% | 6.5% | 6.5% | 6.8% | 5.8% | 5.8% | 6.2% | 5.6% | 5.6% | 5.9% | 10.6% | 6.5% | 7.3% | 10.6% | 12.1% | 14.5% | 15.9% | 22.1% | 25.9% |
| P778 | M | 10w6d | 9.8% | 6.8% | 6.4% | 7.5% | 5.8% | 5.8% | 6.2% | 6.2% | 6.5% | 6.2% | 11.7% | 6.0% | 7.9% | 11.0% | 16.5% | 19.5% | 21.3% | 28.8% | 30.8% |
| P770 | M | 10w5d | 8.9% | 6.9% | 6.3% | 6.8% | 6.4% | 6.1% | 5.8% | 6.9% | 6.1% | 6.8% | 12.5% | 5.7% | 9.7% | 13.9% | 18.8% | 20.6% | 27.2% | 29.4% | 37.3% |
| P679 | M | 10w6d | 13.7% | 7.0% | 6.3% | 7.2% | 7.0% | 6.5% | 6.3% | 6.7% | 6.7% | 6.3% | 12.9% | 6.0% | 8.7% | 14.4% | 16.9% | 21.3% | 25.4% | 29.8% | 28.8% |
| P695 | M | 11w6d | Not available | 7.1% | 6.2% | 6.3% | 7.1% | 6.4% | 6.5% | 6.3% | 6.3% | 6.6% | 12.9% | 6.0% | 8.2% | 12.5% | 17.8% | 21.5% | 26.0% | 28.5% | 34.5% |
| P686 | M | 11w3d | Not available | 7.1% | 7.1% | 6.5% | 7.3% | 6.8% | 6.3% | 6.7% | 6.6% | 6.3% | 11.9% | 5.9% | 7.5% | 12.0% | 19.2% | 18.9% | 26.7% | 26.8% | 42.7% |
| P673 | M | 10w2d | Not available | 7.4% | 6.6% | 7.2% | 7.4% | 7.5% | 7.6% | 6.7% | 7.4% | 6.9% | 12.6% | 5.8% | 9.3% | 14.2% | 20.4% | 20.7% | 21.2% | 27.6% | 29.9% |
| P796 | M | 12w0d | 16.4% | 7.4% | 7.0% | 7.9% | 7.2% | 6.6% | 6.4% | 7.1% | 6.9% | 7.6% | 13.0% | 6.6% | 9.5% | 12.6% | 16.7% | 19.1% | 23.3% | 31.3% | 32.7% |
| P731 | M | 10w1d | 7.4% | 7.5% | 7.3% | 7.2% | 7.5% | 6.7% | 7.0% | 7.0% | 7.0% | 7.3% | 14.4% | 6.2% | 9.1% | 15.5% | 21.0% | 26.3% | 29.3% | 35.7% | 38.6% |
| P693 | M | 11w5d | Not available | 9.1% | 8.5% | 8.4% | 8.8% | 9.1% | 8.7% | 8.9% | 9.1% | 8.7% | 17.0% | 7.9% | 12.0% | 19.1% | 23.8% | 31.1% | 35.1% | 38.2% | 42.8% |
| **Median (n = 28)[e]** | | | **8.9%** | **5.6%** | **5.1%** | **5.8%** | **5.5%** | **5.1%** | **4.9%** | **5.1%** | **5.4%** | **5.1%** | **10.2%** | **4.6%** | **7.3%** | **10.5%** | **13.4%** | **15.5%** | **17.5%** | **23.1%** | **28.4%** |
| **IQR lower, 25th percentile[e]** | | | **6.9%** | **4.9%** | **4.5%** | **4.8%** | **4.5%** | **4.2%** | **4.2%** | **4.2%** | **4.3%** | **4.3%** | **7.7%** | **4.1%** | **5.7%** | **8.2%** | **10.6%** | **12.3%** | **15.0%** | **19.1%** | **24.4%** |
| **IQR upper, 75th percentile[e]** | | | **10.7%** | **6.9%** | **6.4%** | **6.8%** | **6.5%** | **6.2%** | **6.2%** | **6.4%** | **6.3%** | **6.3%** | **12.0%** | **5.9%** | **8.0%** | **12.1%** | **16.7%** | **19.2%** | **22.4%** | **27.8%** | **33.0%** |

**Notes**

a. Marker tiers: the reference of 1,839,680 CCGG sites was ranked by predicted per-CpG fold enrichment and partitioned into octiles of approximately 229,960 markers each, so T1 holds the best-scoring 12.5% of CpGs and T8 the worst. Each library was then restricted to the reads cut at the CpGs of one tier and fetal fraction re-estimated. Median predicted fold enrichment, T8 through T1: 1.1, 2.1, 2.9, 3.5, 4.0, 4.6, 5.3, 6.7.

b. The unfiltered library covers all model markers overlapping in the assay, which is not equivalent to all CCGG sites: 32-38% of read 1 does not fall on the reference at all. It is therefore a different marker set rather than the first step of the tier series.

c. Fetal fraction was estimated from SNPs at which the mother is homozygous using a truncation-corrected maximum-likelihood estimator. Per-case fetal fractions are shown in Supplementary Fig. 6.

d. Tier-restricted epiSIFT values are the apparent fetal fractions within a methylation-selected subpopulation of molecules.

e. Median and interquartile range are computed across patients and describe between-patient spread, not uncertainty on the median. The clinical NIPT column summarizes only the 19 cases with a reported value.
