## Supplemental Figures 1-6 for "Early Prenatal Cell-Free DNA Methylation Landscape Informs Epigenetic Purification of the Circulating Fetal Genome"

### Supplementary Information

#### Table of Contents

**Supplementary Figure 1:** Fragment length distributions of extracted cfDNA from CVS culture media

**Supplementary Figure 2:** Internal consistency of the CVS methylome reference across fetal sex and individual samples

**Supplementary Figure 3:** Methylation concordance between younger and older donors in the normal plasma background reference

**Supplementary Figure 4:** Genomic context and enhancer occupancy of the fetal enrichment marker tiers

**Supplementary Figure 5:** Fetal fraction enrichment by CpG marker tier for patient P352, broken down by chromosome

**Supplementary Figure 6:** Fetal fraction across enrichment marker tiers in individual plasma cases

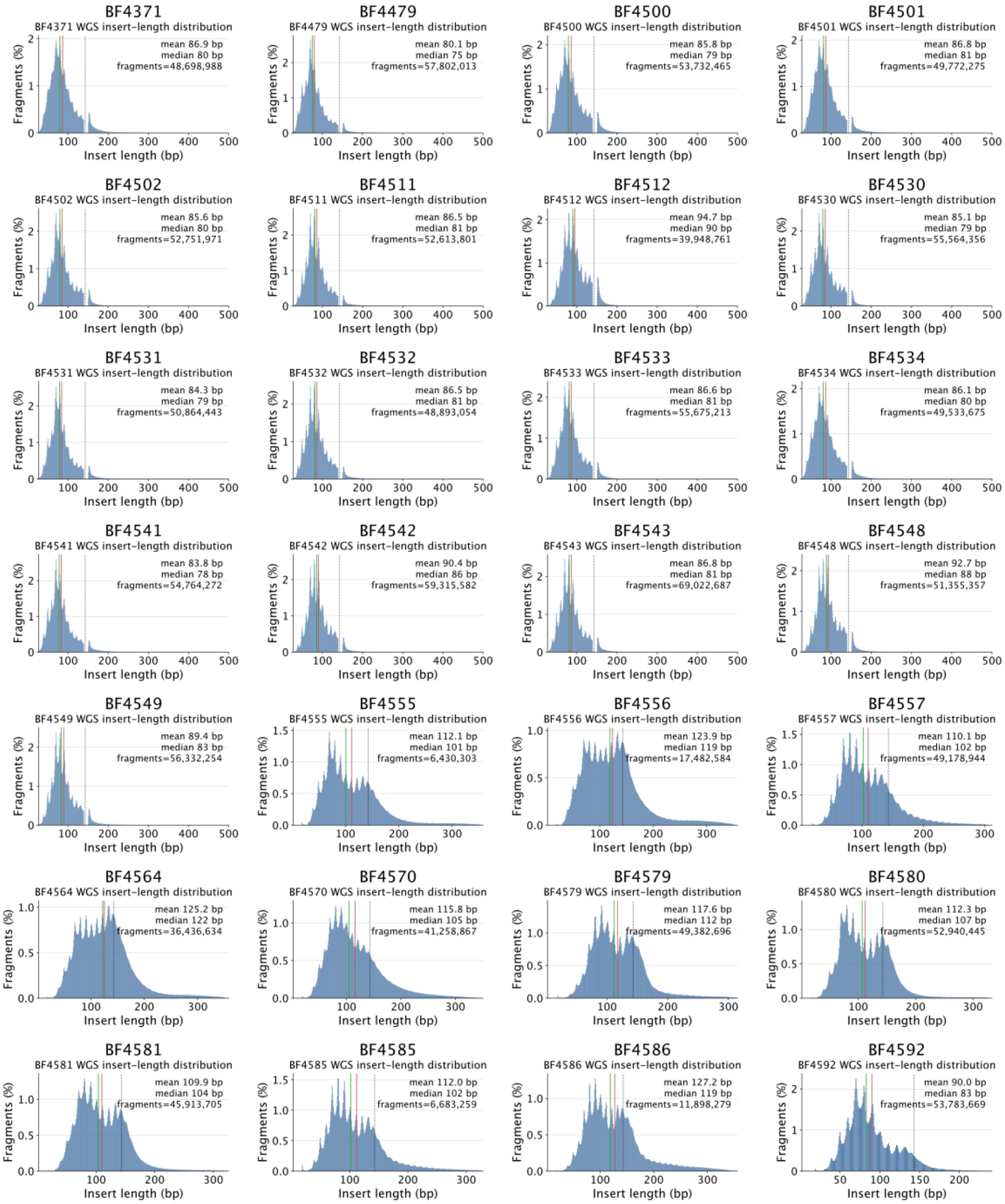


**Supplementary Figure 1: Fragment length distributions of extracted cfDNA from CVS culture media.** Whole genome sequencing length distributions for each of the 28 CVS culture-media samples included in the integrated methylome reference using paired-end insert sizes. Annotations indicate the mean and median insert length and number of fragments analyzed per sample.

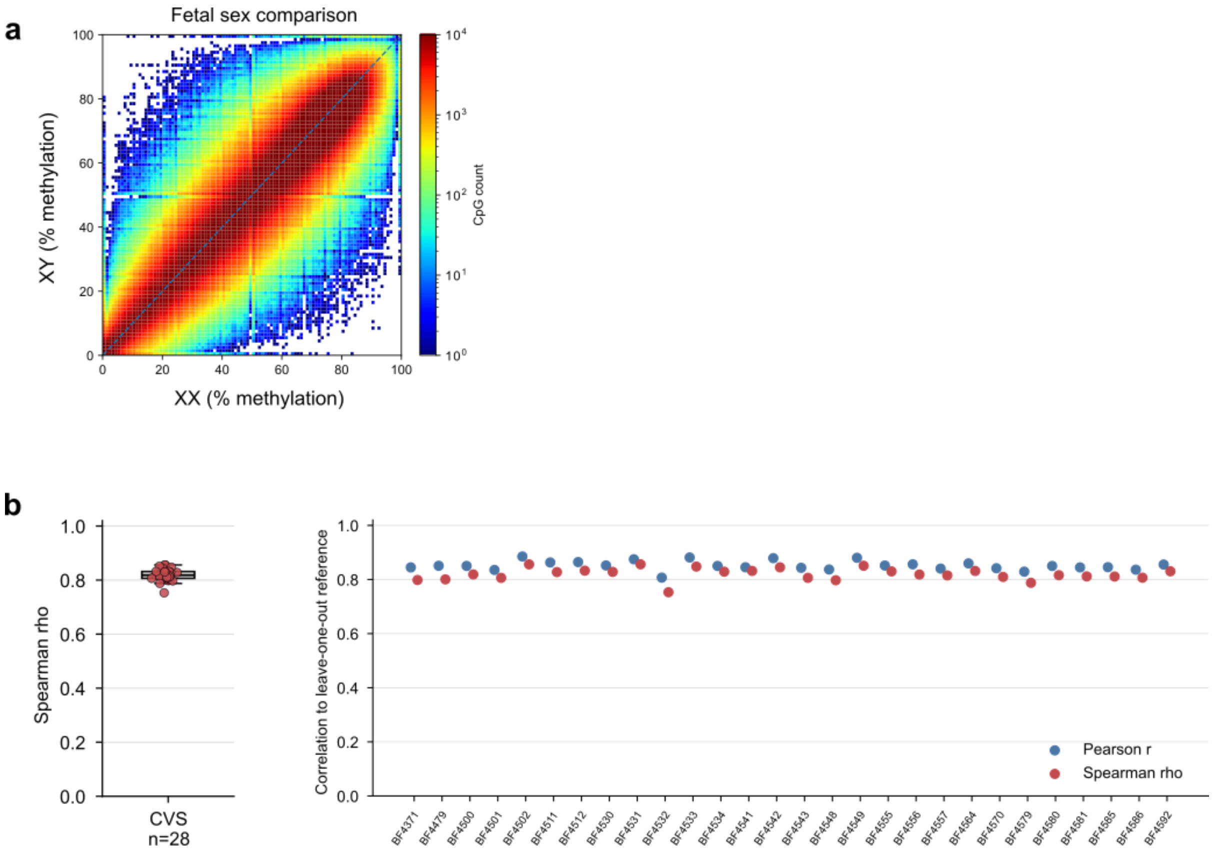


**Supplementary Figure 2: Internal consistency of the CVS methylome reference across fetal sex and individual samples. a,** Heatmap showing CpG methylation concordance (Spearman's ρ=0.936) between female (XX, n=12) and male (XY, n=14) CVS samples. Two XXY samples were excluded from this comparison. **b,** Leave-one-out concordance across the 28 CVS samples in the integrated reference, with each sample compared with a reference constructed from the remaining 27 samples. Left, distribution of Spearman's ρ values with lowest ρ 0.753 and median ρ 0.818. Right, Pearson's r and Spearman's ρ for each sample.

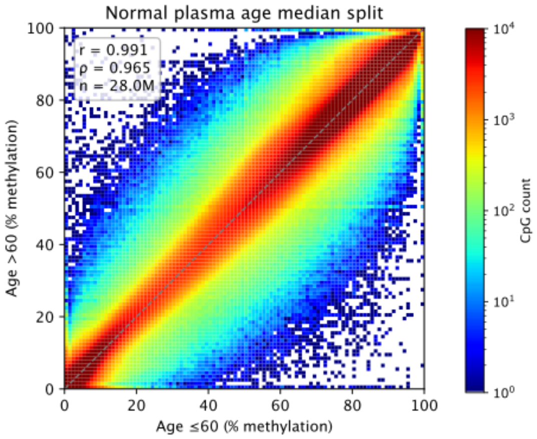


**Supplementary Figure 3: Methylation concordance between younger and older donors in the normal plasma background reference.** Density heatmap of CpG methylation in the normal plasma methylome reference from Loyfer et al. 2023, comparing donors ≤60 years old against >60 years old (Pearson's r=0.991, Spearman's ρ=0.965, n=28.0 million CpGs).

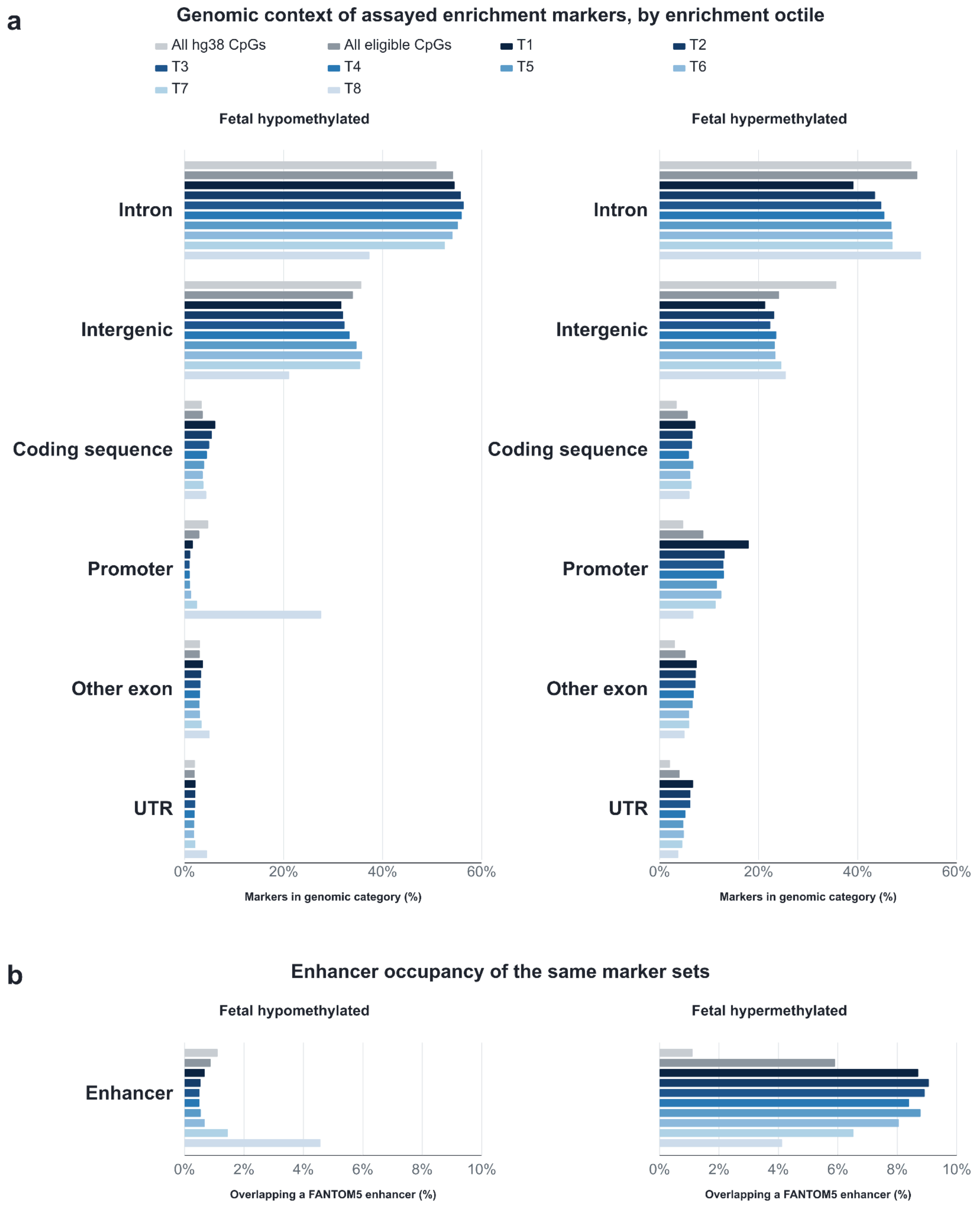


**Supplementary Figure 4: Genomic context and enhancer occupancy of the fetal enrichment marker tiers. a,** Mutually exclusive genomic regions across all hg38 CpGs, all eligible CpGs, and the CCGG-assayable markers in the disjoint tiers of Fig. 3d,e (T1 highest to T8 lowest fold enrichment), separately for hypomethylated and hypermethylated markers. **b,** The same series against non-exclusive, pan-tissue FANTOM5 permissive enhancers.

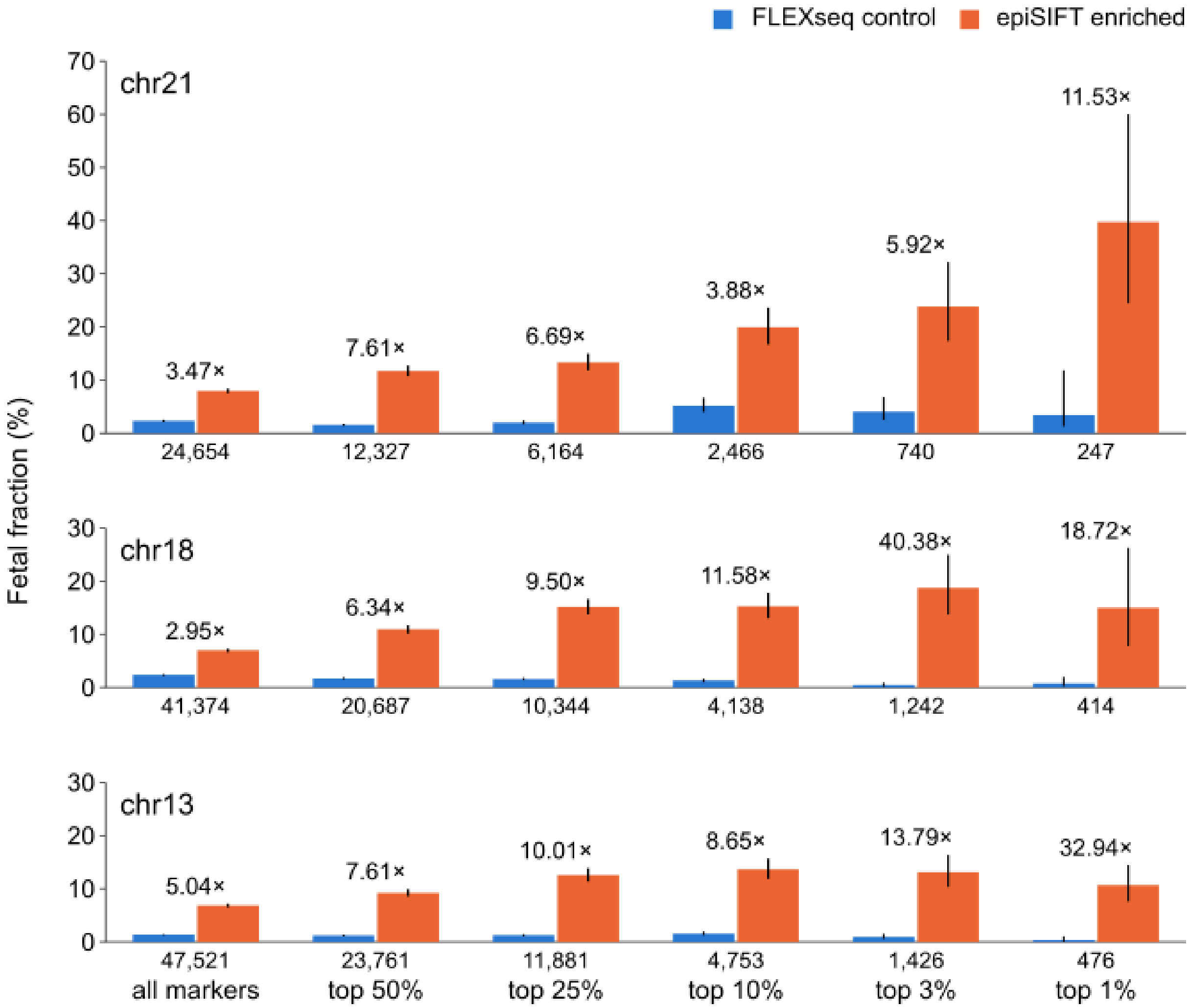


**Supplementary Figure 5: Fetal fraction enrichment by CpG marker tier for patient P352, broken down by chromosome.** Fetal fraction in the FLEXseq control and epiSIFT enriched libraries across all markers and increasingly selective top marker subsets ranked by theoretical fold enrichment, shown for chromosomes 21, 18, and 13.

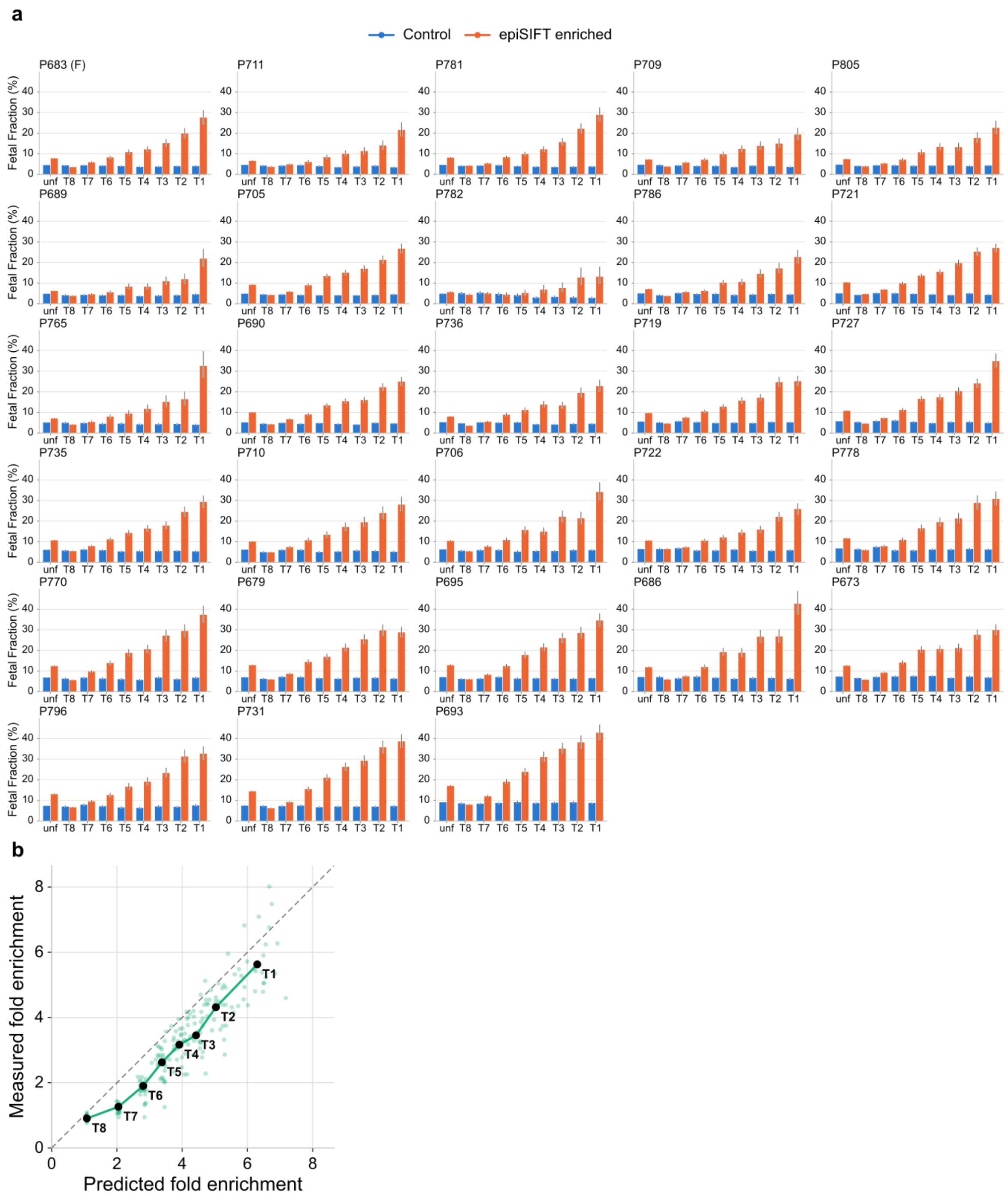


**Supplementary Figure 6: Fetal fraction across enrichment marker tiers in individual plasma cases.** **a,** Fetal fraction in the FLEXseq control and epiSIFT enriched libraries across all overlapping markers and eight marker tiers ordered from lowest (T8) to highest (T1) in theoretical fold enrichment. Each panel represents one plasma case in the cohort of 28. **b,** Measured versus predicted fold enrichment across 8 tiers with the dashed line representing exact agreement. Background green dots are the individual cases with a range of predicted fold enrichment due to variable initial fetal fraction.
